# sCellST: a Multiple Instance Learning approach to predict single-cell gene expression from H&E images using spatial transcriptomics

**DOI:** 10.1101/2024.11.07.622225

**Authors:** Loïc Chadoutaud, Marvin Lerousseau, Daniel Herrero-Saboya, Julian Ostermaier, Jacqueline Fontugne, Emmanuel Barillot, Thomas Walter

**Affiliations:** Centre for Computational Biology (CBIO), Mines Paris, PSL University, 75006 Paris, France; Institut Curie, 75248 Paris Cedex, France; INSERM, U900, 75248 Paris Cedex, France; Institut Curie, Department of Pathology, Saint-Cloud, France; Institut Curie, CNRS, UMR144, Equipe labellisée Ligue Contre le Cancer, PSL Research University, Paris, France; Université Paris-Saclay, UVSQ, Montigny-le-Bretonneux, France; Computational Medicine, Servier Research & Development, Saclay, France

## Abstract

Advancing our understanding of tissue organization and its disruptions in disease remains a key focus in biomedical research. Histological slides stained with Hematoxylin and Eosin (H&E) provide an abundant source of morphological information, while Spatial Transcriptomics (ST) enables detailed, spatially-resolved gene expression (GE) analysis, though at a high cost and with limited clinical accessibility. Predicting GE directly from H&E images using ST as a reference has thus become an attractive objective; however, current patch-based approaches lack single-cell resolution. Here, we present sCellST, a multiple-instance learning model that predicts GE by leveraging cell morphology alone, achieving remarkable predictive accuracy. When tested on a pancreatic ductal adenocarcinoma dataset, sCellST outperformed traditional methods, underscoring the value of basing predictions on single-cell images rather than tissue patches. Additionally, we demonstrate that sCellST can detect subtle morphological differences among cell types by utilizing marker genes in ovarian cancer samples. Our findings suggest that this approach could enable single-cell level GE predictions across large cohorts of H&E-stained slides, providing an innovative means to valorize this abundant resource in biomedical research.

## 1 Introduction

Tissue spatial organization is a fundamental feature of multicellular organisms, essential for proper development, homeostasis, and tissue repair. It depends on the precise arrangement of different cell types and their states, allowing tissues and organs to perform their biological functions efficiently. Disruptions or perturbations in this organization can lead to pathological conditions, such as cancer, where tissue structure and function become compromised.

The most widely used technique to study tissue architecture and its disease related alterations involves Hematoxylin and Eosin (H&E)-stained tissue slides which are routinely produced in clinical practice and examined by pathologists to evaluate disease states and inform treatment decisions. With the advent of machine learning for pathological slides, algorithms have been developed to address tasks such as molecular phenotyping and biomarker discovery[1]. H&E slides offer a detailed view of tissue, capturing both individual cellular phenotypes and overall architecture.

A fundamental step in tissue analysis is the identification of distinct cell types within the tissue. Broad categories, such as epithelial cells, fibroblasts or lymphocytes can be readily identified through manual inspection, due to their characteristic nuclear size, morphology and colour. However, more subtle cell types often require molecular staining, such as Immunohistochemistry (IHC) or Immunofluorescence (IMF), as differential transcriptional programs do not always leave visible morphological fingerprints. In some cases, cell type specific morphological cues may simply remain undiscovered.

More recently, spatial transcriptomic (ST) technologies have emerged as a powerful tool to study gene expression (GE) in the spatial context [2] [3]. These technologies offer complementary analyses to H&E staining by providing insights into the molecular landscape of tissues that are neither accessible through conventional histology nor through IHC or IMF technologies, which are limited in the number of molecular markers. ST technologies can be broadly divided into two main categories. Image-based ST such as MERFISH, COSMIX and Xenium [4] rely on fluorescent imaging and enable the capture of hundreds of different RNA species with highly precise spatial resolution, down to sub-cellular level. However, these methods do not measure the full transcriptome. Furthermore, segmenting cells can be challenging, potentially leading to errors in cell type calling [5]. On the other hand, sequencing-based methods such as Visium, Slide-seq or Stereo-seq use spatial barcodes to retain spatial information at specific locations called spots. Then, next generation sequencing is used to profile mRNA species and reconstruct a spot-gene expression matrix. The downside of these approaches is that spots usually contain several cells, around 10 to 20 in the case of Visium. Deconvolution algorithms [6] [7] are then required to estimate cell type proportions within each spot, with very few of these approaches going back to true single cell level [8, 6]. In addition, both technologies share the downside of being very expensive, which also prevent them from being used in larger cohorts, unlike H&E slides.

Recent studies have demonstrated that H&E-stained slides contain valuable information that can be leveraged to predict GE using machine learning algorithms. One pioneering method in this field is HE2RNA[9], a deep learning model designed to predict GE from bulk RNA sequencing data by aggregating predictions from small patches extracted from corresponding H&E images. Despite the inherent challenges posed by weak and noisy signals, HE2RNA effectively identifies immune cell-enriched regions, suggesting that the link between morphology and GE is sufficiently strong for this kind of approaches. With the advent of ST technologies, particularly Visium, which provides both H&E slides and transcriptomic spot data from the same tissue section, these models have gained even greater predictive power. Two types of approaches have been recently developed: super-resolution models and GE predictors. The former takes image features and spot GE as input to produce super-resolved expression maps [10]. While these models thus virtually increase the resolution inside and between the spots, they still require ST as inputs. The latter refers to models trained to predict GE based on the image centered at each spot locations [11, 12, 13, 14, 15, 16, 17, 18]. Other methods [19], [20] predict GE at finer spatial resolutions using a weakly supervised learning approach. Weak supervision corresponds to a scenario, where the ground truth is not available for each input instance, but only for groups of instances. While these approaches enhance spatial resolution, they still fail to provide estimations of single-cell GE.

Here, we introduce sCellST, a method to predict single-cell GE from cell morphology, trained on paired ST and H&E slides. Our method is versatile and can be applied across different tissues and cancer types, as demonstrated with the diverse datasets used in this study. While designed to make cell-level rather than spot-level predictions, we show that sCellST performs on par with state-of-the-art spot-based predictors on a three-slide pancreatic ductal adenocarcinoma dataset. Next, we validated sCellST’s ability to accurately predict cell types by benchmarking against methods trained on manually annotated images and found remarkable agreement. Finally, we demonstrate that sCellST can help identifying morphological patterns in more subtle cell types by leveraging a list of marker genes extracted from a scRNA-seq dataset.

## 2 Results

### 2.1 sCellST overview

We present sCellST, a weakly supervised learning framework that learns a model to predict GE from H&E images only. Our approach is composed of 3 main steps (Fig.1).

**Figure 1.**
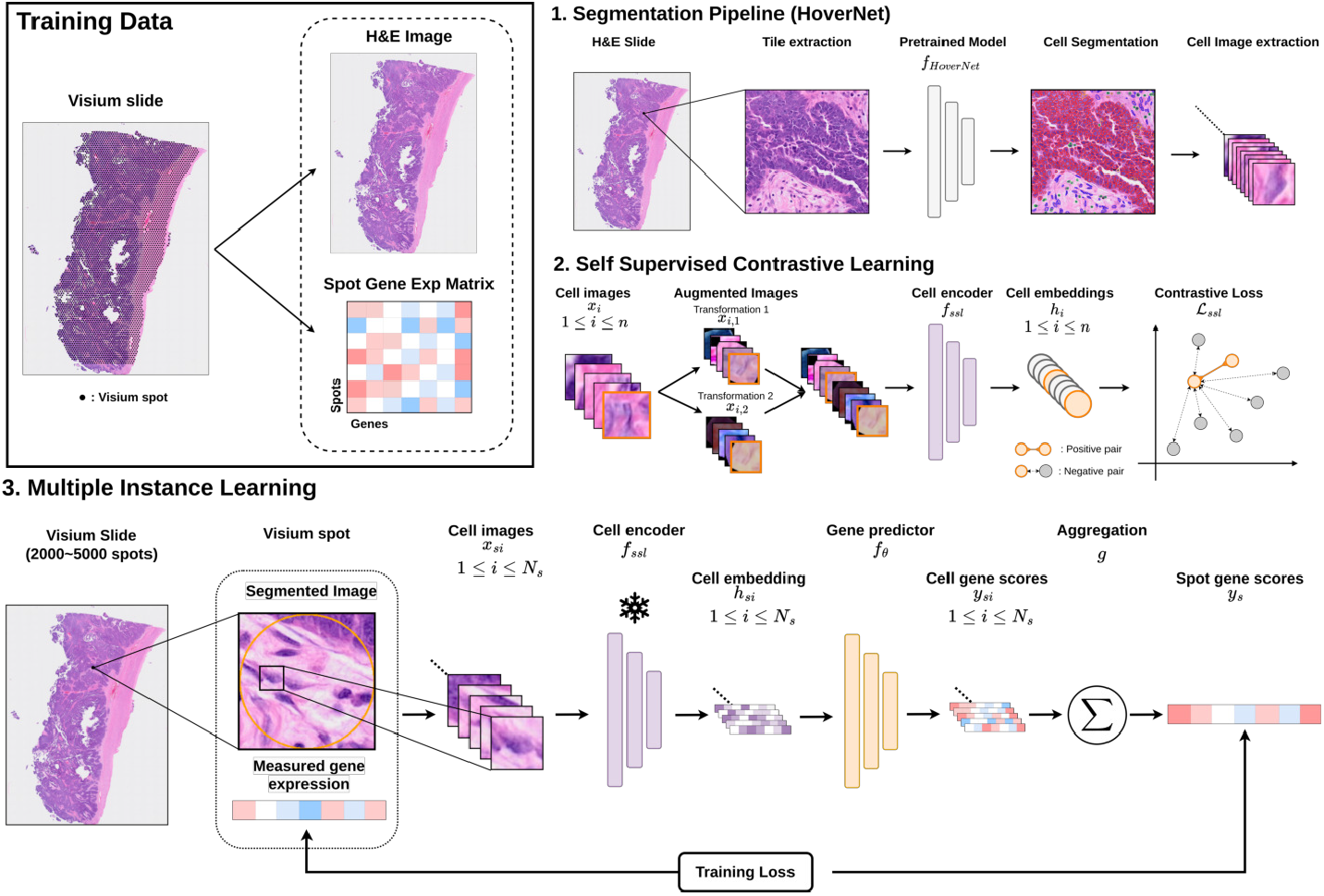
Overview of sCellST pipeline. 1. sCellST first uses a pretrained cell detection algorithm to extract cell images from H&E images. 2. A feature extractor is trained using contrastive learning on the cell images. 3. A Multiple Instance Learning approach is used to learn a GE predictor. The model predicts first GE score for each cell within a spot before aggregating the prediction to produce a spot expression vector which is used for training.

First, a deep learning model is used to perform cell detection on the underlying H&E images from the spatial transcriptomic slide. For each cell detected by the model, we extracted a squared crop images centered on the segmentation output masks. We used HoverNet [21] for our experiments but any other detection algorithm for histopathology data (e.g. [22]) can be used.

The next step is to find a suitable representation of the cell image. For this, we turned to Self-Supervised Learning (SSL), a state-of-the-art method to learn powerful embeddings which can then be used for a large variety of tasks [23]. SSL relies on the definition of pretext tasks, i.e. tasks that are not directly related to the classification problem we want to solve, but for which large datasets are available. It has shown great success on tissue patches and full slides [24], [25] for GE predictions, tile classification or survival predictions. More recently SSL has been used to represent single cell morphologies [26, 27] with great success in distinguishing different cell types. Here, we trained MoCo v3 algorithm [28] (methods).

Finally, we trained a GE predictor using cell images. Since ground-truth GE is not available for individual cells, but rather for spots containing multiple cells, we formulated the problem as a Multiple Instance Learning (MIL) problem [29]. Specifically, for each spot in a Visium slide, we predict a GE vector for each cell within the spot. We then aggregated these predictions to generate a spot-level prediction, which is compared with the measured expression to train the algorithm (methods). The predicted cell-level scores can then be interpreted as cell GE estimates.

After training, sCellST can be applied using only H&E slides to produce single-cell and spatially resolved GE data.

### 2.2 sCellST predicts single cell level gene expression in simulated data

As our datasets do not contain detailed groundtruth on single-cell GE, validation of sCellST at the cellular level is a challenging endeavour. We therefore designed several levels of validation, the first of which is based on simulation experiments using cell images from an ovarian cancer slide.

To establish a ground truth for single-cell GE, we utilized an annotated scRNA-seq dataset as a reference. We matched cell images with GE profiles from this scRNA-seq data according to several scenarios. In the *random* scenario, GE vectors were randomly assigned to cell images, and there was thus no relationship between morphology and GE, providing a random baseline. To artificially introduce a relationship between cell morphology and GE, we first clustered the cell image vectors. Each image cluster was then arbitrarily assigned to one scRNA-seq cluster. In the *centroid* scenario, we assigned to each cell image the mean expression vector of its corresponding scRNA-seq cluster. Finally, we explored a more challenging scenario where we matched cells from the scRNA-seq and from the corresponding morphological cluster (*cell* scenario; see Fig.2a). In this case, as we draw cells randomly from both matched clusters, GE and morphological intra-cluster variation are independent by construction. Once the cell images and GE vectors were matched, we simulated spot-level GE by summing the GE of single cells assigned to each spot (methods).

**Figure 2.**
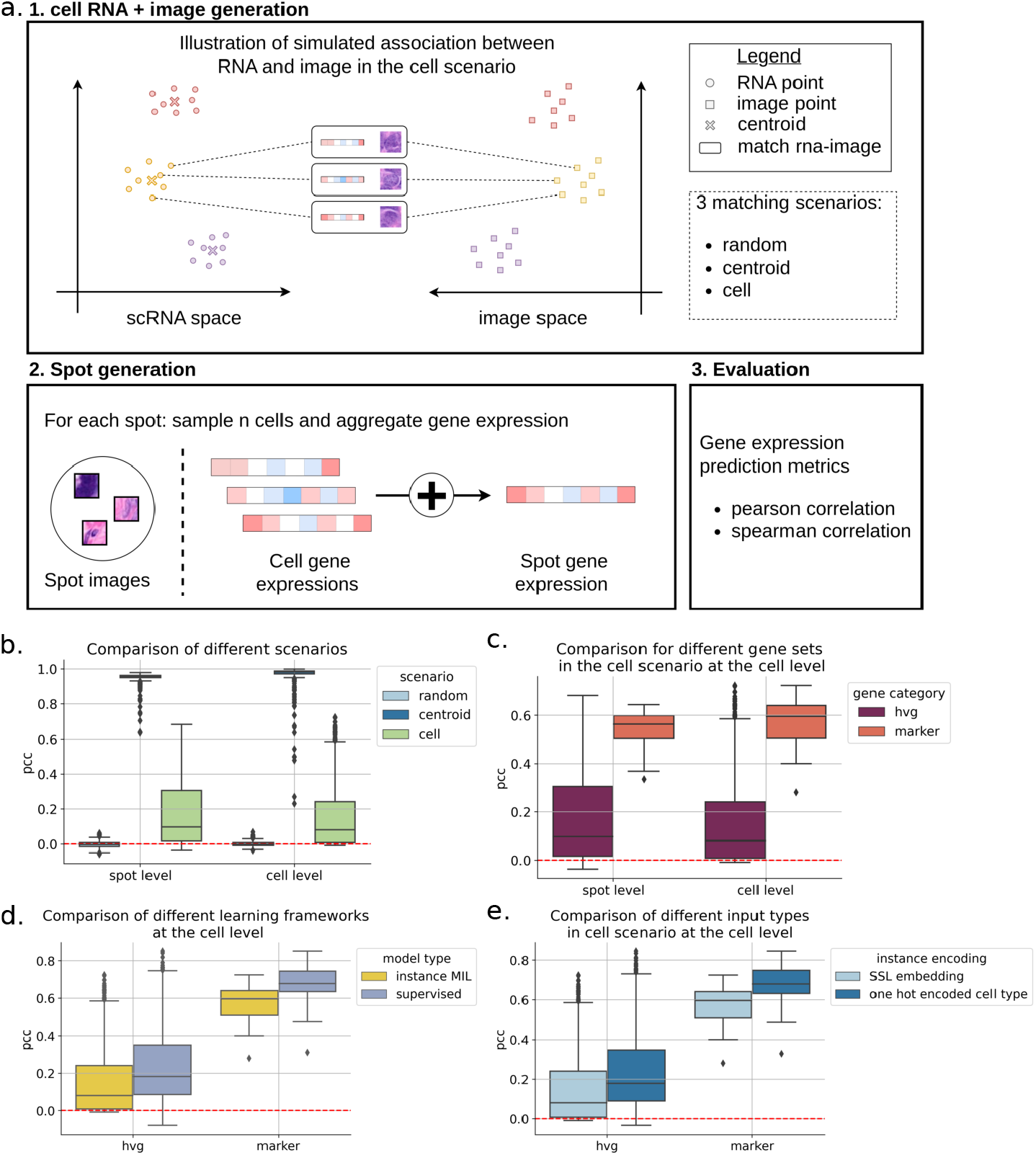
Simulation experiments. a. Simulation framework with cell image - GE attribution with different scenario (cell scenario depicted here). Distribution of Pearson correlations in the test dataset for: b. all genes for the different scenarios, c. different sets of genes, d. different learning frameworks and e. different instance encodings.

We first compared the three scenarios on the top 1000 highly variable genes (HVGs). For both spot and cell-level predictions, genes predicted in the random scenario produced correlations no higher than 0.1, centered around 0 (Fig.2b), as expected in the absence of links between GE and cellular morphology. In contrast, the *centroid* scenario yielded correlations exceeding 0.8 for most genes, demonstrating the effectiveness of the MIL approach.

The *cell* scenario also resulted in positive, non-random correlations. Notably, the marker genes, which are supposed to have stronger links with morphological properties, exhibited an increase in median correlation from 0.10 for HVGs to 0.54 for marker genes (Fig.2c) at the spot level. This indicates that the model can capture varying degrees of correlation between cell morphology and GE.

However, defining an upper bound for the evaluation metric is challenging, as even a model with access to all information would not achieve a correlation of 1, as by design of our simulation, GE is not entirely predictable from morphology. To illustrate this, we trained a model under full supervision to predict the matched GE vectors from the corresponding cell images. The performance of the supervised model was higher than that of the MIL-trained model (Fig.2d), yet still far from reaching perfect correlation (median of 0.18 and 0.68 for HVGs and MG respectively).

Next, we investigated the effect of the tile encoding strategy by replacing the SSL-embeddings by one-hot encoded vectors of the image clusters. These one-hot-vectors represent and ideal noise-free embedding that perfectly represents the cell image clusters. We only observed a mild improvement (Fig.2e), suggesting that the high dimension and intra-cluster variability have only a minor effect on prediction performance.

With these experiments, we demonstrated that a MIL approach can recover GE vectors from cell images, even in challenging settings where ground truth labels for all cell images are unavailable and the link between morphology and GE is only partial.

### 2.3 sCellST performs better than other algorithms on spot level predictions

Next, we aimed to benchmark sCellST against state-of-the-art methods for GE prediction from H&E data. As there is currently no method that specifically targets single-cell GE prediction from Visium data, we performed spot-level comparisons, even though this was not the primary objective of our method. We used 3 pancreatic ductal adenocarcinoma (PDAC) Visium slides (Fig.3b) from [30]. For each slide, we selected common HVGs across all slides which resulted in 321 genes. We evaluated the predictions using Pearson and Spearman correlation coefficients.

**Figure 3.**
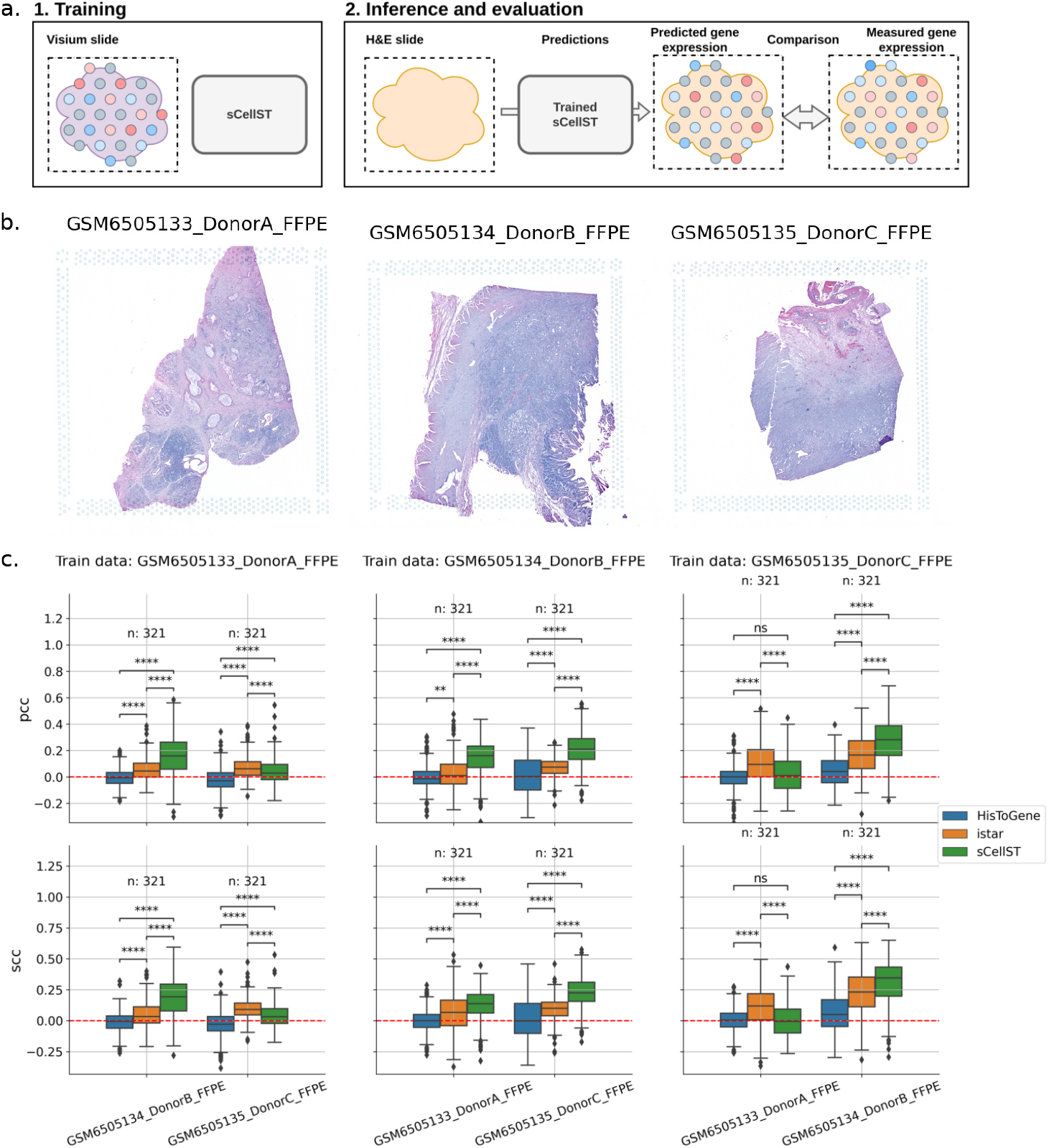
Benchmark of sCellST. a. Overview of the benchmarking approach. Each slide is used for training and then the model is evaluated on the two remaining slides. b. H&E slides from the PDAC Visium dataset. c. Benchmark results: each boxplot represents the distribution of Pearson / Spearman correlation coefficient on all genes.

We compared our algorithm to two methods: HisToGene [18] and Istar [20]. HisToGene is based on a Vision Transformer neural network which takes the image from each spot as input. Istar also utilizes weakly supervised training. It processes small patches from the spot images and predicts GE for each patch, which is then aggregated at the spot level. As shown in our experiments, sCellST outperforms both methods (Fig.3c), which was unexpected given that sCellST was not optimized for this task. To validate the performance difference, we used a Wilcoxon signed-rank test to compare the distributions of results for the HVGs. HisToGene performs poorly, likely due to the large number of model parameters (∼ 300M) relative to the small size of the training dataset (∼ 2000 training spots) compared to Istar and sCellST (∼ 900K). sCellST significantly outperforms Istar in 8 out of 12 experiments (Fig.3c).

We also compared different cell image representations within our model. sCellST using SSL-embeddings are on par with models that utilized embeddings from transfer learning with ImageNet pretraining and outperforms in most cases those which relied solely on one-hot encoded cell type information (SupFig.3), suggesting that more subtle information than what is reflected by the broad cell type categories can be learned.

In most cases, sCellST successfully predicts GE with positive correlations on a set of shared HVGs.

### 2.4 Cell level predictions are consistent with cell type predictions from Neural Networks trained on manual annotations

While our analyses showed that sCellST compares favourably to state-of-the-art spot GE prediction methods, our primary objective was to predict single cell GE. For this, we compared sCellST results with state-of-the-art cell type calling methods trained on manually annotated nuclei. HoverNet [21] is a widely used method for both segmentation and cell type classification, trained on more than 45000 manually annotated nuclei. The main cell types identified by HoverNet are neoplastic epithelial, connective / soft-tissue cells (fibroblasts, muscle and endothelial nuclei) and inflammatory (methods).

In our experiments, we used 2 slides (Fig.4a,b) of breast and ovarian cancer tissue from the 10X Genomics website. For every slide, we applied the sCellST pipeline, restricting the analysis to the top 1000 HVGs. We then grouped cells based on their labels and compared the predicted GE scores between groups. This approach allowed us to identify the top-expressed genes for each cell type label. In Fig.4c,d, we present the top five genes in columns for the three Hover-Net labels. Top differential genes were readily identifiable for the connective and inflammatory cell groups. Regarding connective cells, they encompassed genes involved in muscle contraction (MYLK) and extracellular matrix organization (COL1A2, COL3A1, COL10A1, COL11A1, MMP2), which in turn represent specific markers of stromal cells such as muscle cells and fibroblasts, respectively. For inflammatory cells, we found classical markers of lymphocytes (PTPRC, IGHM), as well as other genes specific to the lymphocytic lineage (LCP1, LSP1, SRGN, POU2AF1). Given the inter-patient transcriptomic heterogeneity of epithelial tumor cells, most differential genes are expected to be patient-specific [31]. Nevertheless, the genes identified are plausible over-expressed genes of the tumoral populations: aberrant over-expression of PTPN3 has been observed in a variety of malignancies, including breast cancer [32], and over-expression of WNT6 and FGF19 is documented in ovarian cancer and linked to several oncogenic mechanisms ([33], [34]). To further validate the predicted GE, we adopted a complementary approach by investigating the predicted expression of established marker genes for each of the 3 HoverNet labels (Fig.4e,f). Classical markers for epithelial tumor cells like EPCAM and E-cadherin (CDH1) showed higher levels in cells labelled as neoplastic, as did canonical inflammatory myeloid cells (LYZ, CD68) and fibroblasts (COL1A2, INHBA).

**Figure 4.**
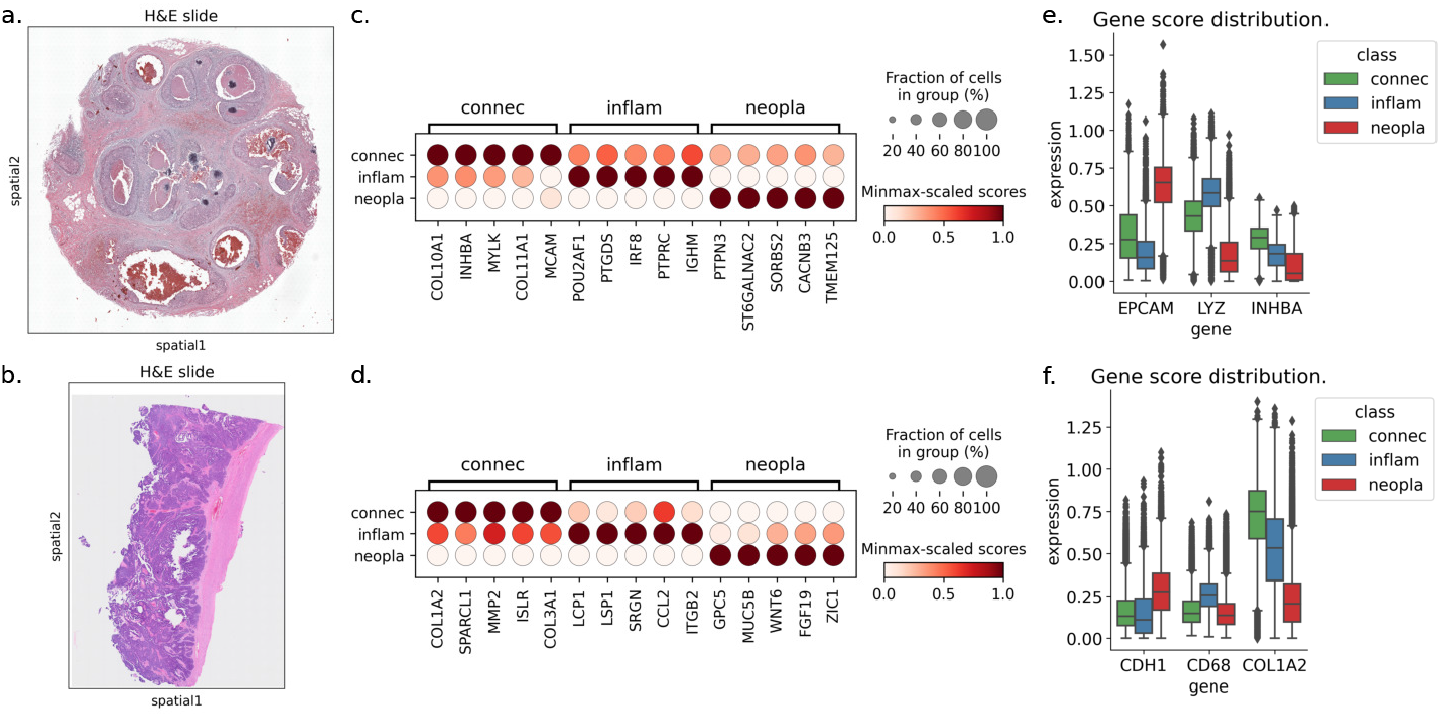
sCellST comparison with HoverNet labels: Each row corresponds to a Visium slide from cancer tissue: a, c, e: breast and b, d, f ovarian. a,b. Visium slides used for the experiments. c,d: Top differentially expressed genes when grouping cells by HoverNet labels. e,f: Distribution of known marker genes grouped with HoverNet labels.

Next, we visually examined the coherence of HoverNet and sCellST predictions. For each of the 2 Visium slides, we show a crop of the H&E image, the cell segmentation with cell types predicted by HoverNet, the spots with Visium measurements and the segmented cells coloured with sCellST predictions (Fig.5). The single-cell GE predictions provide fine-grained information on the cell-type, in line with HoverNet classification results, yet more detailed. In the two crops shown, sCellST allowed us to detect the real pattern of organization of immune cells, consisting of densely populated clusters at the edges of the tumor mass. This pattern was impossible to detect with Visium resolution. In the breast cancer slide, we observed a thin layer of connective cells encapsulating the tumor that exhibited high predicted expression of INHBA. These fibroblasts could correspond to the ECM-myoCAFs described by [35] or the INHBA+ CAFs described by [36]. In the ovarian cancer slide, the Visium spots showed high expression of the myeloid marker CD68 in the tumor region; however, we did not observe an accumulation of immune cells at the corresponding locations. Conversely, sCellST allowed us to assign CD68 expression to the immune cells found in the interstitial stromal regions outside of the tumor, a pattern usually recognized as immune-excluded, with important implications in the clinic [37].

These analyses provide qualitative evidence of our model’s relevance by showing that the distribution of predicted GE is consistent with established biological knowledge.

**Figure 5.**
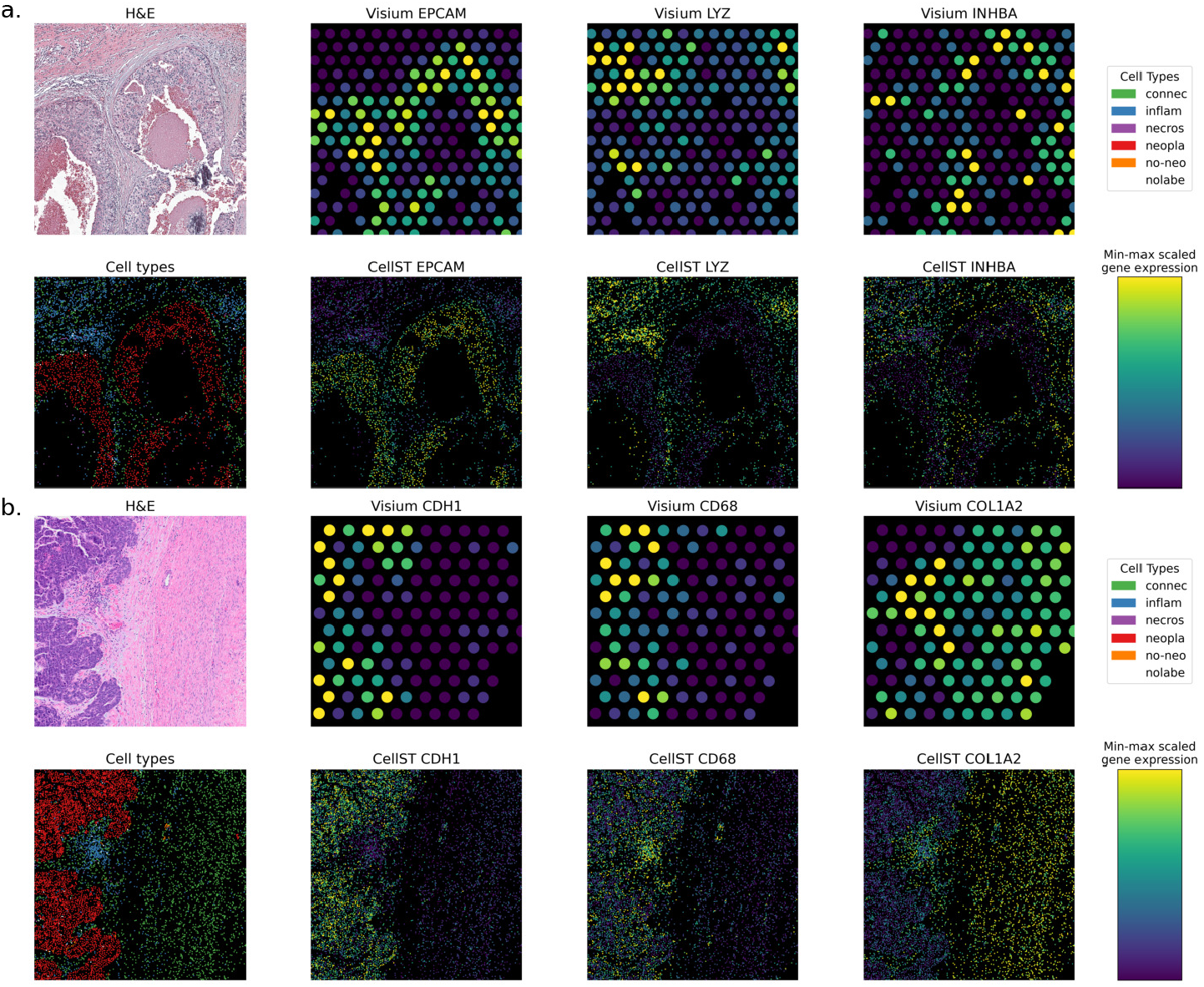
Cell level predictions with sCellST. Each subpanel corresponds to a Visium slide from cancer tissue: a: breast and b: ovarian. For all slides, the first row corresponds to the H&E image followed by Visium measurement in spots for 3 genes, the second to cell types predicted by HoverNet and then sCellST gene prediction for the same genes.

### 2.5 sCellST can identify finer cell types in ovarian cancer

Broad cell type labels obtained with algorithms trained with cell type annotations can be limiting, as multiple finer cell types may be grouped into a single annotated category. In this study, we show how marker gene lists can be used to identify the morphology of more specific cell types. We focused on a publicly available human ovarian cancer slide from the 10X website. We trained our model to predict a list of marker genes which were obtained from an annotated single-cell dataset of ovarian tissue available on the CellXGene website [38]. After training the model, we used single-cell GE predictions to define cell type scores for each cell (methods). We computed cell type scores for five cell types: fibroblasts, endothelial cells, lymphocytes, plasma cells, and fallopian tube secretory epithelial cells. We then compared sCellST scores with the broad categories provided by HoverNet. Cells labelled as connective showed higher sCellST scores for fibroblasts and endothelial cells, while those labelled as inflammatory had elevated lymphocyte scores in sCellST. Lastly, cells classified as neoplastic exhibited high scores for the epithelial type in sCellST.

To understand the model’s ability to distinguish finer cellular subtypes based on morphology, we subsequently generated cell image galleries by plotting the 100 cells with the highest score for each cell type (Fig.6c). For connective tissue cells, including fibroblasts and endothelial cells, distinct morphological characteristics could be identified from the top-scoring cell images. Although both fibroblasts and endothelial cells were spindle shaped, endothelial cells tended to be less elongated and to line a vascular space, sometimes containing red blood cells, further corroborating their identity. For inflammatory cells, the lymphocyte image gallery revealed small round cells with dark nuclei and scant cytoplasms. In contrast, plasma cells were larger and ovoid, with more abundant cytoplasms and eccentric nuclei. Additionally, we noted potential misclassification by HoverNet, as cells with high plasma cell scores—labelled by HoverNet as either connective or neoplastic—exhibited morphological characteristics previously associated with plasma cells (SupFig.5). Of note, while the overall spot performance of SSL-and ImageNet-encodings is similar, galleries obtained with ImageNet-encodings are less homogeneous for plasma cells and endothelial cells (SupFig.4).

**Figure 6.**
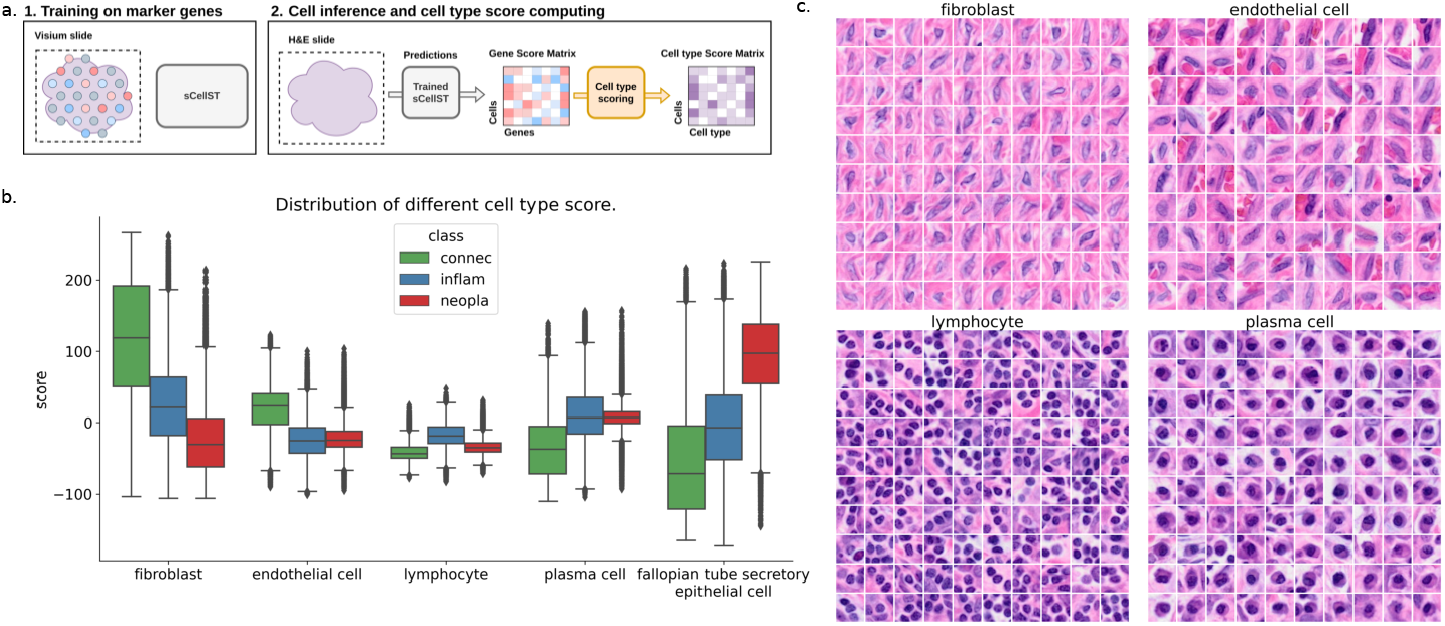
sCellST discovers cell type morphological features. a. Schema of the approach. First sCellST is trained on a set of marker genes obtained from prior knowledge. Then, a scoring function is used to produce cell type scores for each cell. b. Distribution of scores for several cell types grouped by HoverNet labels. c. Correlation heatmap of cell type scores. d. Image galleries with highest score images for each cell type.

In conclusion, we have demonstrated that our model, when integrated with cell type marker genes, can effectively identify cells displaying distinct morphological characteristics that are not captured by state-of-the-art cell type classification models. For instance, our analyses suggest that sCellST succeeds in distinguishing fibroblasts and endothelial cells, while such subtle categorization is currently not available with state-of-the-art cell type classification models. Importantly, this analysis was conducted without reliance on manually annotated labels, yet the model successfully identified known morphological patterns solely using the predicted GE scores.

### 2.6 Discussion

We presented sCellST, a method for predicting single-cell and spatially resolved GE from H&E images based on a weakly supervised learning framework. Unlike other approaches, sCellST generates a detailed spatial map of cell-type-specific expression patterns from H&E data, providing a more granular understanding of GE at the single-cell level.

Although not originally designed for this purpose, we demonstrated that sCellST achieves either superior or comparable performance to state-of-the-art models for the spot level GE prediction task. Furthermore, we showed that sCellST provides results in line with state-of-the-art methods trained on tens of thousands of manually annotated nuclei on broad cell type categories. Importantly and in contrast to such methods, sCellST can also identify more finegrained cell types relying on subtle morphological differences which would be difficult or impossible to generate manually.

For these reasons, we believe sCellST has the potential to drive several important developments. First, it enables large-scale studies of the relationship between nuclear morphologies and GE, facilitating the identification of cell type-specific morphologies. Second, sCellST introduces a novel annotation strategy for single-cell computational pathology (SCCP), which currently depends heavily on extensive manual annotations. For example, Diao et al. trained cell classification models on 1.4 million manually annotated cells by certified pathologists [39]. sCellST offers an efficient way to generate large-scale single-cell annotations with minimal manual input, while also distinguishing more fine-grained cell types. As such, sCellST could significantly impact SCCP. Finally, the ability to dissect cell types from H&E images opens up unprecedented opportunities to reanalyse existing H&E cohorts for predicting outcomes and treatment responses. Although ST is a powerful technique, it is unlikely to be applied to large retrospective cohorts in the near future due to its high cost and limited tissue availability. sCellST offers the possibility to create high-resolution virtual ST. While virtual ST may not fully match the quality of direct measurements, it is still expected to provide valuable and exciting insights.

The proposed method is not free of limitations. First, sCellST relies on cell segmentation, making it vulnerable to segmentation errors, which can become more pronounced with differences in staining and scanning. These technical variations also affect the generalizability of the cell image embeddings, ultimately limiting the broader use of our model without retraining. Future research could address this by developing domain-robust cell image representations. Added to this, ST data is still scarce and consequently, the small size of training datasets negatively impacts predictive performance and generalisation. Some initiative starts to collect massive amount of data such as [40, 41] but the availability of FFPE Visium slides and access to the corresponding high-resolution images remain limited. At the same time, spatial resolution of sequence-based ST is increasing [42]. In Visium HD, bins are arranged on a 2 *μ*m rectangular grid. This higher resolution could enhance GE prediction but requires custom training techniques, as bins are typically grouped into 8 μm sizes and do not align with individual cells.

Overall, we believe that our approach can serve as a pioneering method for predicting GE from cell morphology in H&E images. With the scaling of ST dataset sizes, it has the potential to predict GE on large cohorts of H&E images, facilitating novel biological discoveries.

## 3 Online Methods

### Notations

- *S* number of spots in a Visium slide
- *G* number of genes
- *f*_*θ*_ a neural network parametrised by a set of weights *θ*
- *x* a cell image
- *h* a cell embedding vector
- *y* ∈ ℕ^*G*^ a raw vector of gene expression and *Y* ∈ ℕ^*S*×*G*^ the spot GE matrix
- *y*^*p*^ ∈ ℝ^*G*^ a preprocessed vector of GE and *Y* ^*p*^ ∈ ℝ^*S*×*G*^ the preprocessed spot gene expression matrix
- We used the notation *ŷ* to denote predictions, in this case of a raw GE vector.
- *p*_*NB*_(.; *a, b*) density function of a negative binomial distribution parametrised by the parameters *a* and *b* corresponding to either *μ* and *θ* used in [43] or total counts and probability of success parametrisation.

### Spatial Transcriptomics datasets

We based our approach on Visium technology because it provides both a spatially resolved transcriptomic profile and a corresponding H&E image. Visium is part of the spot-based ST family, capturing mRNA using spatial barcodes within defined spots that typically contain 10–20 cells. The captured mRNA is then sequenced using next-generation sequencing (NGS) technology. A key advantage of this method, compared to image-based ST, is its ability to capture the entire transcriptome, though at a lower spatial resolution. In our study, we utilized Formalin-Fixed Paraffin-Embedded (FFPE) ST datasets, as FFPE preserves cellular morphology more effectively than fresh frozen tissue samples. Additionally, we analysed H&E images at the highest available resolutions, ranging from 0.2 to 0.5 *μ*m per pixel, depending on the specific slide.

### ST preprocessing

For each ST dataset, we filtered genes with less than 200 counts and those that were detected in less than 10% of the training spots. Furthermore, we filtered spots for which no cell was detected and those with less than 20 counts. These filtering steps exclude genes with very low expression on the slide and therefore unlikely to be predictable, as well as uninformative spots. In our experiments, we used either custom gene lists (based on marker genes known to be informative on cell types) or Highly Variable Genes (HVGs) selected on the training spots. For the benchmarking studies, we used 2000 HVGs in order to have sufficient overlap between the three slides. For other experiments, we used 1000 HVGs.

### Cell segmentation from whole slide images

For the segmentation step, we utilized a publicly available pre-trained network called HoverNet [21], which simultaneously performs cell segmentation and classification. We employed the implementation available on GitHub (https://github.com/vqdang/hover_net) and used pre-trained weights from the PanNuke dataset, enabling classification into six main cell types For each segmented nucleus, we extracted a 12*μ*m × 12*μ*m (typical cell size) image centered on the cell’s segmentation center coordinates, which was resized to a 48 × 48 pixel images. Based on the spatial coordinates of the spots, cells, and the spot radius, we linked each cell to its corresponding spot. This kind of models is usually not applied to the whole slide for memory reasons but to tiles (small patches) which cover the tissue. We replaced the mask algorithm from HoverNet used to detect tissue with the one from https://github.com/trislaz/Democratizing_WSI/blob/main/src/tile_slide.py since we find it to work better for the slides we used. Briefly, it combines an Otsu algorithm with a morphological opening to compute the mask. The algorithm used in this study, HoverNet, predicts six distinct classes: neoplastic epithelial cells, connective/soft-tissue cells (including fibroblasts, muscle, and endothelial nuclei), inflammatory cells, non-neoplastic epithelial cells, dead cells, and unlabelled cells. In our analysis of the H&E slides, we observed that the majority of cells were classified into one of the first three categories. Therefore, we restricted our analysis to these three primary classes.

### Image embedding

Given the limited amount of data available for training the GE predictor, we employed strategies to obtain image embeddings independently from this prediction task. Specifically, we explored two approaches: Transfer Learning from ImageNet classification and Self-Supervised Learning (SSL). In both cases, we used a ResNet-50 backbone as encoder. In both scenarios, NN are utilized to map input images to generic representations. For transfer learning, these representations are obtained by training the NN on entirely different data and entirely different tasks (such as classification of natural images), while SSL is optimized for the same type of data (in our case histopathology data), but is trained on so-called pretext tasks, that do not require any annotation or external groundtruth. Among the various available SSL methods, we opted to use MoCo v3, a contrastive learning framework. Briefly, this approach generates two different views of each input image (i.e., transformed using specific augmentations) and is optimized to pull together the corresponding representations for views originating from the same image, while pushing apart those that originate from different images. Choosing the right augmentations for image transformation is key to obtain good representations and can heavily influence the performances of downstream tasks [44]. Since H&E images are very different from natural images, we changed the augmentations and worked with different parameters for colour jittering. We also added two other transformations: random erasing and random rotations. For the latter, we extracted larger cell images as to avoid rotation induced artifacts. To train the SSL network, we used all cells from the training slide which might results in lower performance than expected in cross validation experiments because of a relatively low number of images for SSL training (≤ 300K) but it corresponds to best practice in the field to avoid data leakage.

In two experimental settings, we used one-hot cell type encodings as cell image representations. This corresponds to a vector that is one for the correct cell type, and 0 otherwise.

### Multiple Instance Learning (MIL) for spatial transcriptomics

Unlike the classical supervised learning framework, Multiple Instance Learning is designed for learning in the case when a single label *y* ∈ 𝒴 is available for a set of instances {*x*_1_, *x*_2_, …, *x*_*k*_}, referred to as a bag. Although each instance in the bag has an underlying label, these individual labels remain inaccessible during training.

In the instance-level approach of MIL algorithms, the objective is to learn an instance-level predictor *f*_*θ*_ which assigns a score to each instance. These instance-level scores are then aggregated using a function *g* to generate a score for the entire bag. As the order of instances in the bag is irrelevant, *g* must be permutation invariant. Common choices for *g* include mean or max operations, depending on the specific task.

As every operation presented above can be chosen differentiable, the instance-level predictor *f*_*θ*_ is often parametrised using neural networks, which has proven effective, particularly when working with image data. Such approaches are frequently applied to tasks in computational pathology in the general formulation of MIL [45].

The instance-based MIL framework is particularly well-suited for spot-based ST, as each spot *s* within a slide can be seen as a set of cells which represent the instances. For each spot index *s* in 1, …, *S*, we have a target GE vector *y*_*s*_ ∈ ℝ^*G*^ and a set of images 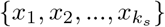 derived from the detection algorithm where *k*_*s*_ represents the number of cells in the spot *s*.

Cell embeddings 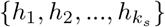 are produced using a pretrained embedding model *f*_*φ*_.

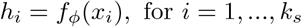

A feed-forward neural network, *f*_*θ*_, is then trained to predict a vector of GE scores based on the cell embeddings. The goal was to develop an algorithm which could predict biologically relevant GE scores at the single-cell level. In order to ensure positiveness of the single cell GE scores, we used a softplus function as final activation function. Finally, a mean aggregation function was applied to simulate the measurement process and derive a GE estimate for comparison with the measured spot-level GE.

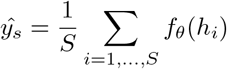

### Model optimization

Two types of objective functions were considered for training the GE predictor.

The first utilized a mean squared error (MSE) loss with preprocessed GE data. In this approach, GE values were normalized by library size and logtransformed, following standard practices in spatial transcriptomic and singlecell analyses:

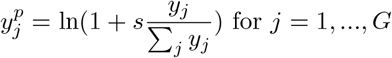

*s* is a normalisation constant set to 10000. In this case, the loss function becomes:

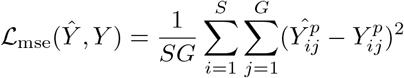

To make sure that all genes contributes to the loss, we additionally scaled each gene between 0 and 1.

The second objective function was based on minimising the negative log-likelihood on raw counts by modelling GE data using a negative binomial distribution. In this formulation, predictions were made for either the mean or the total count parameter of the distribution. The other parameter *α*_*j*_, chosen to be gene specific, is learned during the training. In both cases, the observed library size *l*_*i*_ was used as a scaling factor to avoid capturing this information in the single-cell scores. In this case the loss function becomes:

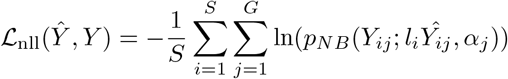

Both approaches were evaluated through simulations; no significant advantage was observed with the negative binomial formulation, as illustrated in the simulation experiments (SupFig.2). Thus, the MSE-based approach was retained for subsequent analyses.

### Simulations

We used a scRNA-seq dataset from CellXGene of ovarian cancer cells and an H&E slide from a Visium slide to perform our simulation study. After training the SSL model on cells extracted from the H&E, we clustered the image embeddings with a k-means algorithm (k=20) to identify distinct morphological clusters from which we kept 6 clusters after visual inspections. We kept only the closest 2000 cells to each cluster centroid in order to have strong morphological differences between clusters. To assign GE vectors to each cell image, we matched the morphological clusters with annotated clusters from the single-cell RNA-seq dataset. GE was then assigned to each cell image based on three scenarios to evaluate the model’s performance under different levels of association between GE and cell morphology:

1. Centroid scenario - perfect link between cell morphology and GE: The mean GE of the corresponding scRNA-seq cluster was assigned to each cell image (SupFig.6.a)
2. Random scenario - no link between cell morphology and GE: Each cell image was assigned a random GE vector from the scRNA-seq dataset (SupFig.6.b)
3. Cell scenario - partial link between cell morphology and GE: Each cell image was assigned the GE vector of a cell from the corresponding scRNA cluster (SupFig.6.c)

For each scenario, we generated 5,000 spots, with each spot containing 20 cells for both training and testing sets. These numbers were chosen to reflect the setting typically observed in Visium ST slides. The model was trained using the top 1000 highly variable genes (HVGs) selected on the spots from the cell scenario training set. To evaluate model performances, we computed the Pearson and Spearman correlations, at both the spot and cell levels, on log-normalized GE vectors for models trained with mean squared error and on normalized GE vectors for models trained with negative log likelihood.

We present comparisons with Pearson correlation coefficient in the main text and with Spearman correlation coefficient in SupFig.1.

### Cell prediction downstream analysis

Following cell GE prediction using a trained sCellST model, we utilized standard tools in Scanpy [46] for downstream analysis. Specifically, we performed differential expression analysis using the “rank genes groups” function of Scanpy with a t-test to rank genes after grouping cells based on HoverNet labels. For the cell type analysis with marker genes, we first identified cell type marker genes with a reference single-cell dataset [47] from CellXGene. We selected only cells originating from the left and right ovary. We applied a t-test to identify 20 marker genes per annotated cell type cluster. B and T cells were merged to form a lymphocyte group, and we excluded mast cells, monocytes, and dendritic cells from the analysis because of the difficulty to unambiguously recognize them in H&E images and thus to validate the morphology galleries produced by our method. From the scRNAseq dataset, we identified 98 marker genes (see Sup. Table 1), of which 93 remained after filtering (see above). We subsequently used the “score genes” function of Scanpy to compute signature scores for cell type marker genes. For each cell *i*, the scoring process involved two lists: a marker gene list *G*_*m*_ and a control gene list *G*_*c*_. First, we normalized the predicted GE values, then the score of a cell for the marker gene list *G*_*m*_ was calculated as follows:

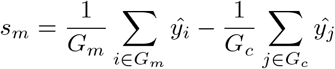

### GE predictors

We compared our approach to two other state-of-the-art methods for GE prediction from H&E data. We made some modifications in the original code to adapt them to the training data presented in this study and to make fair comparisons.

- HisToGene: HisToGene is based on a Visual Transformer architecture which take as input images of spots alongside binned spatial coordinates. As it has been originally implemented for Spatial Transcriptomics data (previous version of Visium) which contain fewer and larger spots per slide, we increased the number of positional encodings from 64 to 128 to enable error-free model training.
- Istar: Istar is a weakly supervised approach that trains five neural networks and aggregates their predictions to obtain the final output, a technique known as *ensembling* in machine learning and statistics. For this study, we reduced the number of trained models from five to one to ensure fair comparison with HisToGene and sCellST. Indeed, ensembling can be applied to every method and usually enhances performance.

## 4 Data Availability

The data are publicly available and links to download raw data are provided below:

spatial transcriptomic slides:

- ovarian cancer: https://www.10xgenomics.com/datasets/human-ovaq rian-cancer-11-mm-capture-area-ffpe-2-standard
- breast cancer: https://www.10xgenomics.com/datasets/human-breas t-cancer-ductal-carcinoma-in-situ-invasive-carcinoma-ffpe-1 -standard-1-3-0
- pancreatic ductal carcinoma dataset: https://www.ncbi.nlm.nih.gov/geo/query/acc.cgi?acc=GSE211895

single cell RNA dataset: https://cellxgene.cziscience.com/e/b252b01 5-b488-4d5c-b16e-968c13e48a2c.cxg/

## 5 Code Availability

The code for this manuscript will be publicly available on GitHub at https://github.com/loicchadoutaud/sCellST.

## 6 Acknowledgements

This work was funded by the French government under the management of Agence Nationale de la Recherche as part of the “Investissements d’avenir” program, reference ANR-19-P3IA-0001 (PRAIRIE 3IA Institute). Furthermore, this work was supported by ITMO Cancer (20CM107-00).

We would like to thank Lucie Gaspard-Boulinc, Nicolas Captier, Nicolas Servant and Loredana Martignetti for helpful discussions.

## 7 Author contributions

L.C, M.L., E.B, and T.W. designed and planned the study. L.C., M.L. and J.O developed the tool. L.C., D.H. and J.F. performed the analysis. L.C, D.H., J.F., E.B. and T.W. wrote the manuscript. All authors reviewed and/or edited the manuscript before submission.

