## Supplementary Material for "sCellST: a Multiple Instance Learning approach to predict single-cell gene expression from H&E images using spatial transcriptomics"

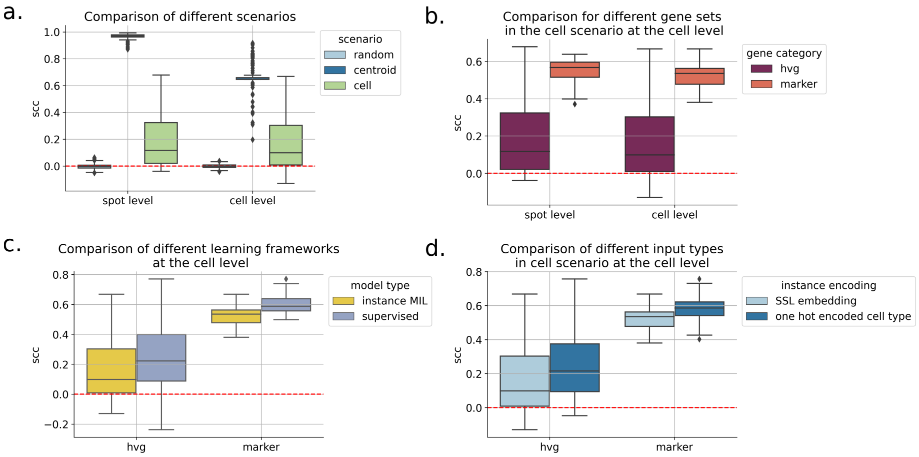


Figure 1: Evaluation with Spearman correlation. a. Distribution of correlations for all genes in the test dataset for the different scenarios with the log1p normalised approached approach. b. Comparison of performance when considering different sets of genes. c. Boxplots showing the differences between different gene expression modelisation.

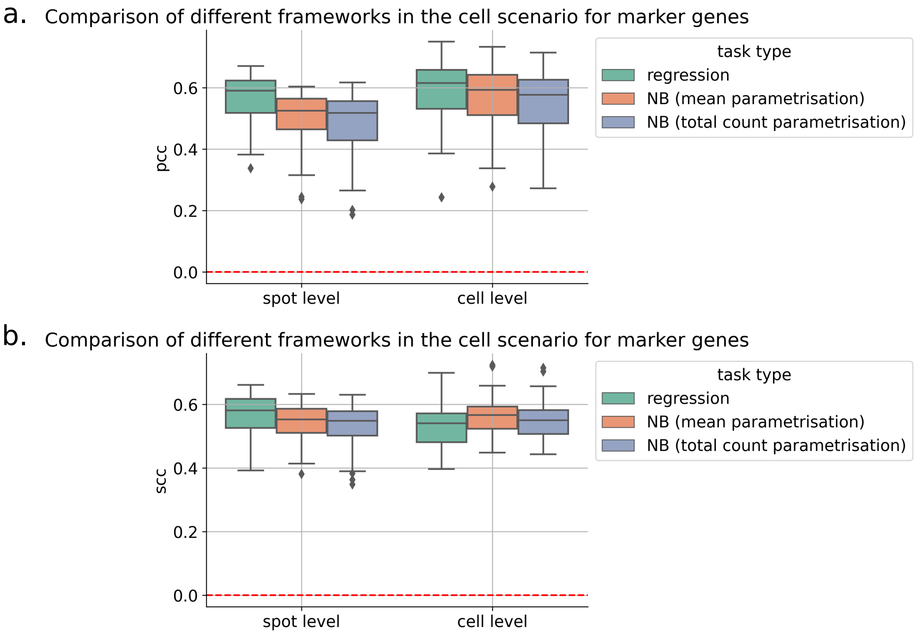


Figure 2: Comparison of performances when using different parametrisation of the loss function.

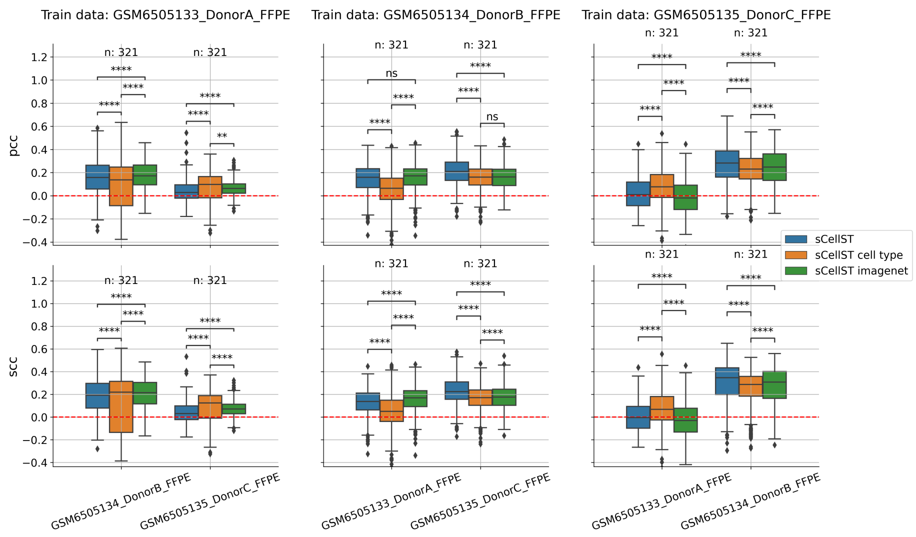


Figure 3: Benchmark results: each boxplot represents the distribution of Pearson / Spearman correlation coefficient of all evaluated genes for different input types.

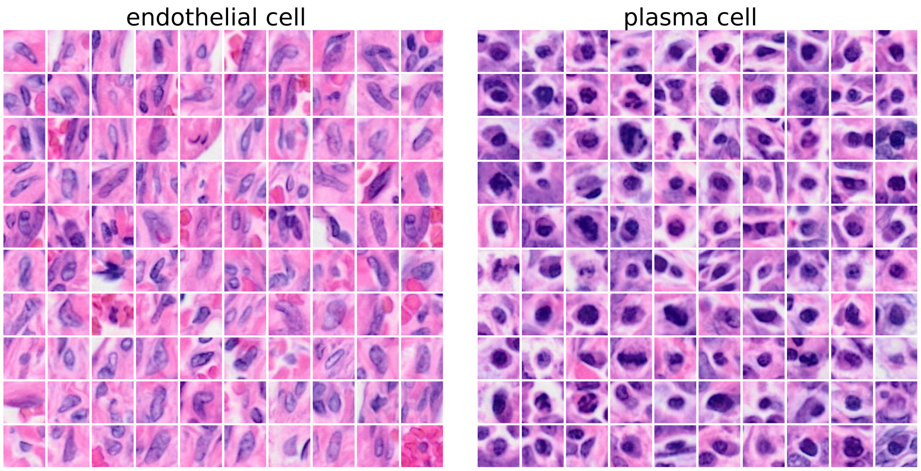


Figure 4: Image galleries with highest score images for each cell type using ImageNet embeddings

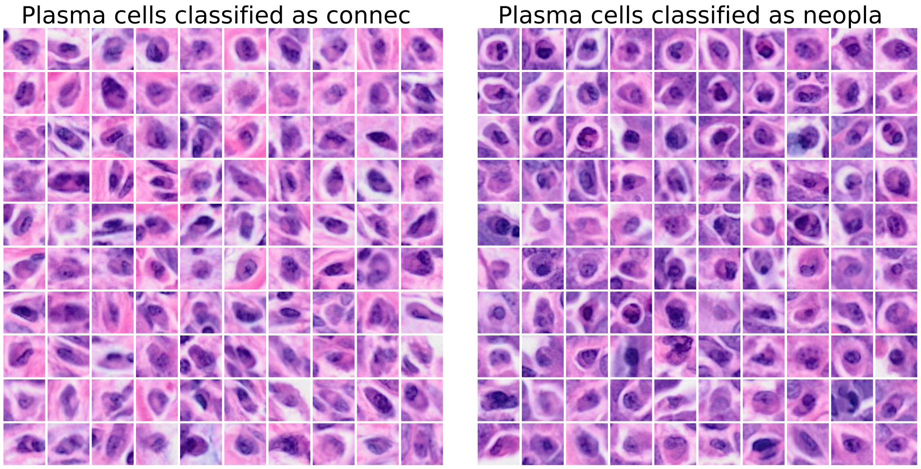


Figure 5: Image galleries with highest score images for plasma cells classified as connective or neoplastic by HoverNet.

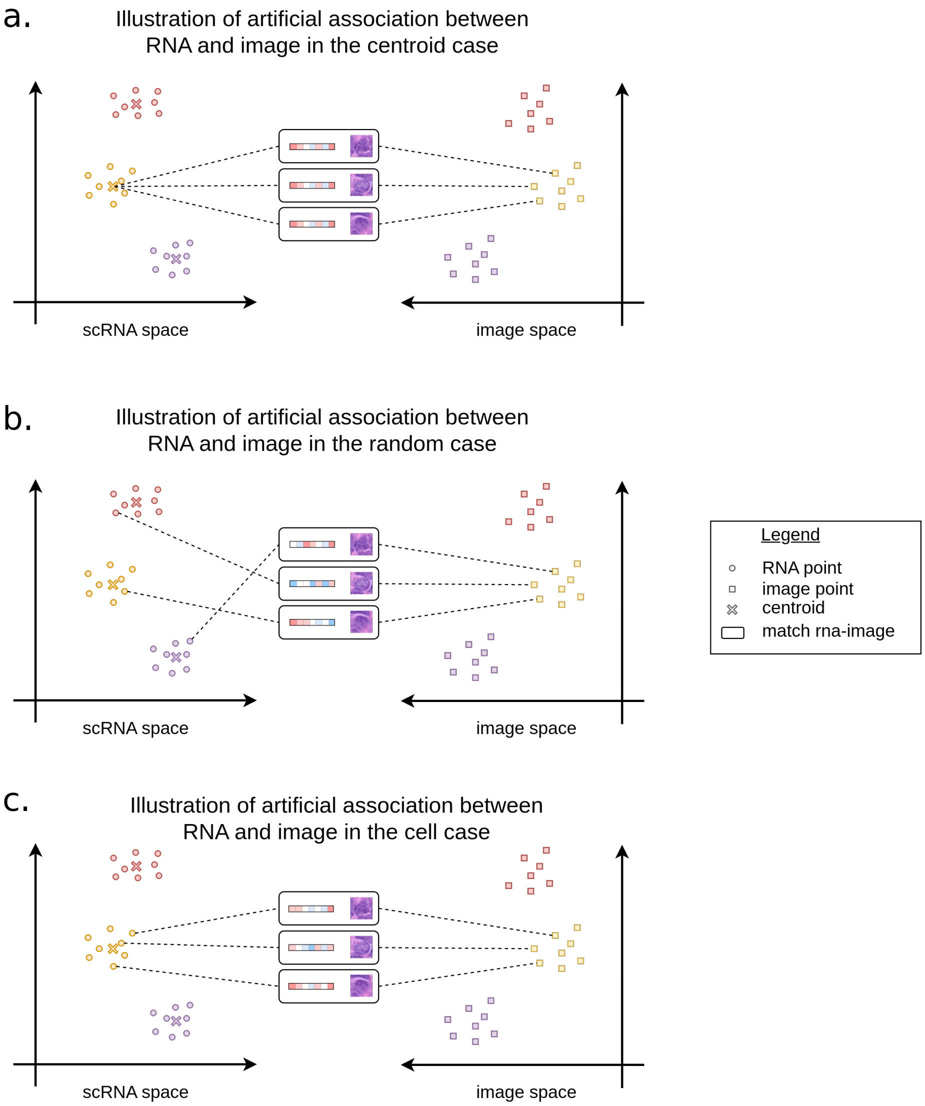


Figure 6: Different scenario for the simulation process of gene expression attribution to cell images.

Table 1: Marker genes extracted from the ovarian scRNA-seq dataset.

| cell type | top 20 genes |
|---|---|
| fibroblast | [AEBP1, C1R, C1S, CALD1, COL1A1, COL1A2, COL3A1, COL5A1, COL5A2, COL6A1, COL6A2, COL6A3, CTHRC1, DCN, LGALS1, LUM, PCOLCE, RARRES2, SPARC, VIM] |
| endothelial cell | [A2M, ADGRL4, APP, CD34, CD93, CLEC14A, COL4A1, COL4A2, EGFL7, ENG, GNG11, HSPG2, IGFBP7, PECAM1, RAMP2, SPARC, SPARCL1, SPTBN1, VIM, VWF] |
| lymphocyte | [B2M, BTG1, CCL5, CD2, CD3D, CD3E, CD52, CORO1A, CXCR4, ETS1, EVL, GZMA, HCST, IL32, NKG7, PTPRC, TMSB4X, TRAC, TSC22D3, ZFP36L2] |
| plasma cell | [CD79A, DERL3, FCRL5, FKBP11, FKBP2, HERPUD1, IGHG1, IGHG3, IGHG4, IGKC, ITM2C, JCHAIN, MZB1, PRDX4, SEC11C, SPCS3, SSR3, SSR4, TENT5C, XBP1] |
| fallopian tube secretory epithelial cell | [BCAM, CD24, CD9, CLDN3, DSP, ELF3, EPCAM, KRT18, KRT19, KRT7, KRT8, LAPTM4B, MSLN, MUC1, RPL35A, RPL8, S100A13, SLPI, SPINT2, WFDC2] |

Table 2: Number of cell per class detected by HoverNet in H&E slides

| | class |
|---|---|
| connec | 62521 |
| inflam | 25887 |
| necros | 3707 |
| neopla | 252712 |
| no-neo | 4638 |
| nolabe | 2790 |

(a) Ovarian cancer slide

| | class |
|---|---|
| connec | 14092 |
| inflam | 16539 |
| necros | 1428 |
| neopla | 12696 |
| no-neo | 88 |
| nolabe | 361 |

(b) Breast cancer slide
